# From static to dynamic: how object rotation influences grasp decisions in ambiguous settings

**DOI:** 10.1101/2023.11.20.567846

**Authors:** Natalie J Maffitt, Emma G Cairns, Julia Wozna, Demetris S Soteropoulos, Alexander Kraskov

## Abstract

The hand grasp of an object is normally consistent, determined by the optimal, most efficient strategy learnt from previous experience. Yet in certain settings, despite object properties remaining constant and intention the same, the grasp chosen by an individual can vary with a lack of clear preference for a particular grasp configuration. This is referred to as motor ambiguity. Here, we compare the influence of preceding static versus dynamic visual information on participants’ choice between two possible hand configurations when grasping an object at various orientations. We confirm previous findings that presentation of an object in an initial static orientation prior to the grasping in ambiguous orientation causes subjects to select a grasp that would be most congruent with the initial, determinate orientation. However, we unexpectedly found that when object rotation is observed between the initial and target positions, the bias is inversed, i.e. subjects choose the alternate grasp configurations. Furthermore, the inverse bias strength was found to be independent of the motion magnitude, and persists even when greater decision time is allowed. We provide evidence that this bias effect is not a perceptual phenomenon, and suggest it is potentially a behavioural manifestation of inhibitory circuits that delay decisions in conditions of motor ambiguity.

## Introduction

On any given day humans perform hundreds of movements that involve interacting with our environment. Any motor action requiring contact with an object requires consideration of its appearance, weight, location, and context in order to successfully complete the task according to need (Rosenbaum et al., 2012). From even the most basic of actions such as grasping a pencil to the more complex skills that take years to develop (e.g. learning to play the violin), each task can present with several different ways to completion. The response selection for a given task can be consistent, determined by the optimal, most efficient strategy learnt from previous experience. Yet in some cases, despite object properties remaining constant and intention the same, the action chosen is variable. The term coined to refer to a lack of clear preference in response choice is ‘motor ambiguity’ (Wood and Goodale, 2011).

Take for example the familiar action of reaching out to pick up a pen placed on a table perpendicular to the body’s sagittal plane. When grasping with the right hand, it comes naturally to position the right thumb on the side of the pen proximal to the body, and the index finger on the distal side. As the pen rotates clockwise on the table to approach alignment with the body’s sagittal plane, it continues to be natural to grasp in the same configuration as before (visually the right thumb is now placed on the pen’s left side), yet it is also reasonable, and more importantly comfortable, to grasp by placing the right index finger on the pen’s left side and right thumb on the right by rotating the forearm. This principle of motor ambiguity can also be extended to hand choice and situations wherein an object can be grasped using a precision or power grip (Seegelke et al., 2016).

A movement towards an always identical object can vary in execution depending on the context presented before movement initiation. This phenomenon has been investigated since the 1990s when Craighero and colleagues reported that participants are quicker to initiate movement towards a target if they had previously been presented with a priming object congruent in orientation with the target (Craighero, 1998, Craighero et al., 1996). In the context of motor ambiguity, the width of the previously presented object can influence an individual’s grip aperture when moving to reach for an ambiguously sized object (Garrido-Vásquez et al., 2021). Likewise, the grip orientation used to grasp a circular object can be influenced by the orientation of a previously seen elliptical object (Hesse et al., 2008). This has been described by some as ‘proactive influence’, i.e. the movement preparation associated with the first object can influence the motor plan for a secondary target (Gallivan et al., 2016). In these examples, the preceding information can be described as static and unchanging, briefly shown for a moment as a technique to prime/influence the motor planning network prior to grasp execution. Yet, it is common in real-life situations for the information available prior to grasping to behave dynamically, i.e., it can change over time, for example preparing to catch a falling object before a gust of wind changes its trajectory. In this study we aimed to explore how the motor planning network is influenced as prior information transforms from static to dynamic, and the consequences this has for behaviour in ambiguous settings.

There have been previous theoretical proposals for how motor planning may be influenced by prior information. The real-time motor control view refers to the idea that it might be more efficient to generate a single movement plan only when the action is required, rather than to continually update the plan as the object changes position, sometimes referred to as ‘dorsal amnesia’ and part of the two-visual systems hypothesis (Westwood and Goodale, 2003, Schenk and Hesse, 2018, Goodale and Milner, 1992). Thus, one of our predictions (see below) was that motor planning would not engage until object motion had ceased, thus preventing proactive influence (Prediction 2).

We first sought to confirm the findings of Hesse et al., 2008, and Gallivan et al., 2016, with the prediction that an object orientation shown prior to grasping influences the grasp orientation chosen by the individual. To achieve this, we designed a grasping task with a target object that, depending on orientation, placed subjects into decision settings where grasp selection was either clear and consistent (determinate), or uncertain and variable (ambiguous). This grasping target was presented after subjects were first exposed to the same object positioned at either a determinate or ambiguous orientation, allowing us to induce the proactive influence effect. Consistent with previous literature, we confirmed that the visual presentation of a static determinate initial orientation prior to the grasping of an ambiguous target orientation biases subjects to select a grasp for the ambiguous target that would be most congruent with the initial, determinate orientation. In a subsequent exploratory experiment, we revealed object rotation between the first ‘priming’ orientation, which is observed, and the second ‘target’ orientation, with three alternative possible predictions.

Prediction 1: Persistence of proactive influence in spite of object motion, i.e. the grasp that would be used for the priming orientation is subsequently used for the target orientation (bias effect).

Prediction 2: Object motion prevents proactive influence, leading to no influence of the priming orientation on the grasp used at the target orientation (no bias effect). This would be in line with the theoretical views regarding dorsal amnesia.

Prediction 3: Object motion inverses proactive influence, i.e. the opposite grasp to that which would be used for the priming orientation is used at the target orientation (inverse bias effect).

After finding evidence in support of Prediction 3, we further hypothesised that this inverse bias effect would diminish with how much time subjects were given to observe the target orientation post-rotation before grasping, and investigated the possible underlying mechanisms by exploring subjects’ visual perception of the task. Our findings provide insights into the interaction between mechanisms of influencing motor planning and its consequences for grasping in motor ambiguity settings.

## Materials and Methods

Ethical approval for all experiments was granted by the Newcastle University Ethics Committee and written informed consent was obtained from all volunteers. This study was performed in accordance with the guidelines established in the Declaration of Helsinki, except that the study was not preregistered in a database.

### Grasping Studies

#### Participants

All participants except for two were right-handed and all were naïve to each experiment and its predictions. A total of 20 healthy volunteers (22-50 years old, 13 females 7 males) were recruited to take part in different experiments in this study (Static Exp.: 10 participants; Dynamic Exp. 1: 12 participants; Dynamic Exp. 2: 10 participants; Dynamic Exp. 3: 10 participants). Some participants took part in several experiments - in which case each experiment was separated by at least 6 days.

#### Grasping Studies - Task Instrumentation

In total, four experiments (Static Exp. and Dynamic Exp. 1-3) were conducted to investigate the influence of prior visual information on grasp selection in ambiguous settings. The experimental paradigm required participants to perform a precision grip movement on a regular triangular prism block (6 cm long, 2 cm base side, see Fig. 1) with their right-hand. Participants were seated in a chair with their forearm positioned on the arm rest (Figure 1a). A circular ‘home-pad’ button (6 cm in diameter) was placed at the distal end of the arm rest and used as the starting location for the wrist. The target object was positioned 35 cm anterior to the centre of the home pad and 15 cm superior, with its rotation occurring in the frontal plane with the rotation axis pointing towards the subject. This resulted in the object lying 25 cm to the right of the body’s midline. As comfort and therefore choice of grasp has been shown to be sensitive to body and head posture (Wood and Goodale, 2011), a chin rest was used to mitigate any postural shifting that could introduce unwanted variability to behaviour. The target object was designed with the intention to afford only one possible grasp: the thumb and finger at opposing ends of the prism block (Stelmach et al., 1994). The object was mounted onto a grey rectangular stage and connected to a stepper motor to enable rotation in steps of 0.9° angles – the stepper motor was driven by an Arduino UNO R3 and motor shield R3, and controlled by custom written Arduino scripts. Thin triangular metal plates were glued to each end of the prism block and connected to a custom made circuit that allowed detection of skin contact by a change in impedance. By placing a glove onto the index finger to impede electrical contact with the object, grasp configuration selection could be recorded in terms of thumb placement, either ‘Grasp 1’ whereby the thumb was in contact with metal plate 1 or ‘Grasp 2’ whereby the thumb was in contact with metal plate 2 (Figure 1a). Previously studies have referred to these two grasp postures as ‘thumb-left’/’supination’ or ‘thumb-right’/’pronation’ respectively (e.g. Stelmach et al., 1994; Wood and Goodale, 2011), however given both of these nomenclatures become uncertain at certain object/wrist orientations, we opted for operational definitions: Grasp 1 and Grasp 2, as defined above.

**Figure 1.**
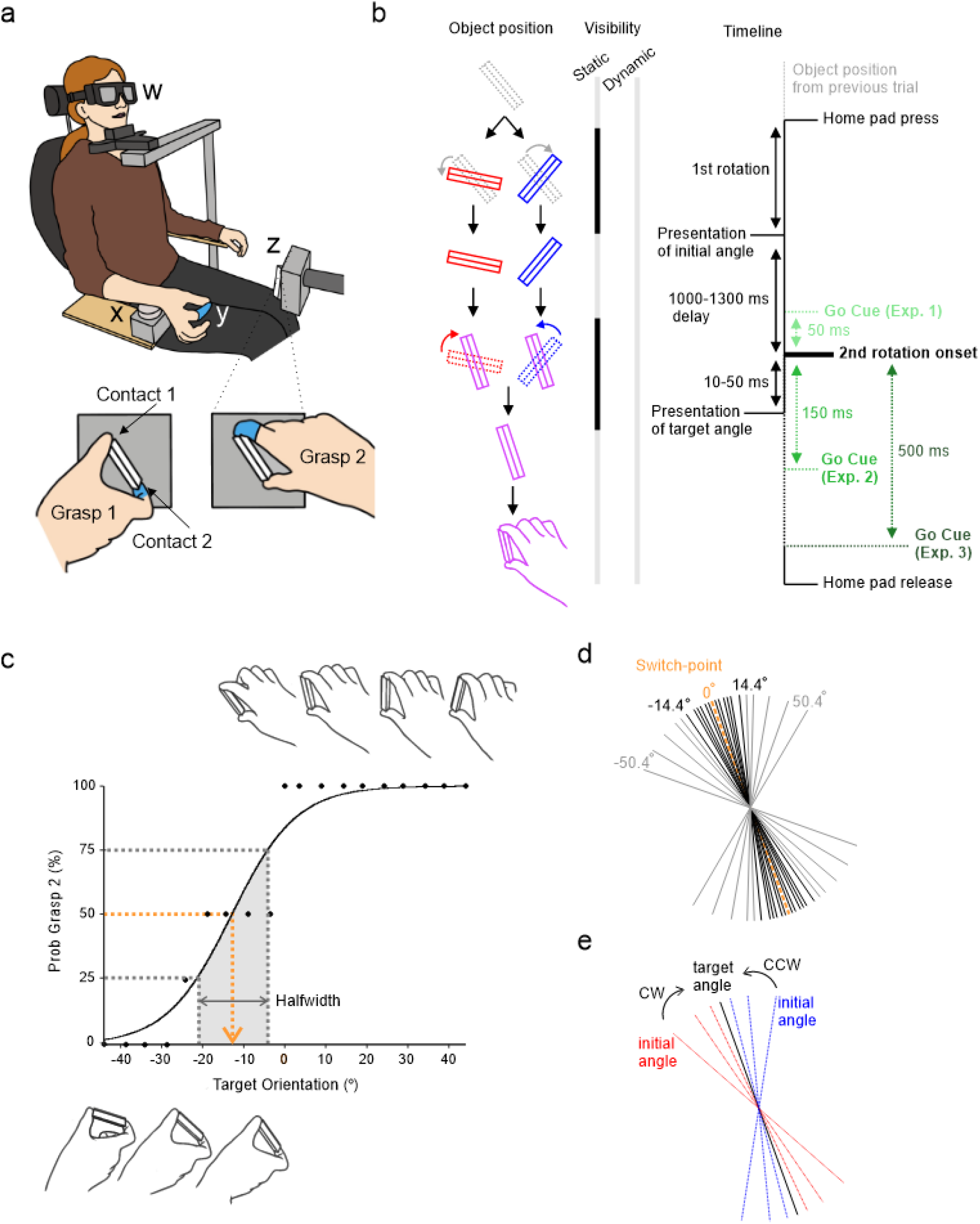
Illustration of experimental task set-up and study design. a) Participant is seated with LCD goggles (w) covering their visual field and head placed on a chin rest. Their right-wrist is used to press down the home-pad button (x), initiating the trial sequence. A latex glove on the index finger (y) impedes electrical connection with the contact circuit on the target object (z) to allow trial-by-trial identification of grasp choice (Grasp 1 or Grasp 2). b) Comparative experimental timeline. The duration of go cue was 50 ms, with variable onset relative to object rotation onset from initial to target angle depending on experimental condition (Dynamic Exp. 1 – -50 ms, go cue starting 50 ms before object rotation and ending at rotation onset, Dynamic Exp. 2, 4 – 150 ms and Dynamic Exp. 3 – 500 ms after rotation onset; time of go cue is indicated by vertical black arrows pointing downward). Periods of object visibility and obscurity are illustrated by light and dark bars respectively. c) Black dots show probability of choosing a Grasp 2 posture for each target angle sampled. Illustrations of object position at example target angles are shown above and below the graph. Estimation of the midpoint was done by taking the 50 ^th^ percentile (orange dotted line) on a logistic fit of the probability of using a Grasp 2 posture in the Calibration task. Estimation of transition sharpness (‘Halfwidth’) between the two grasp choices is measured as the range of angles for which the probability of Grasp 2 lies between 25% and 75% (grey dotted lines). d) Trial design of Experimental tasks showing sampling of target angles, with ‘0°’ representing the subject’s switch-point. In total 23 angles, symmetric around 0°, spanning -50.4° to +50.4° range with denser sampling of ambiguous (close to switch point) were used (list of all target angles -50.4, -39.6, -29.7, -21.6, - 18.0, -14.4, -9.9, -8.1, -6.3, -3.6, -1.8, 0.0, 1.8, 3.6, 6.3, 8.1, 9.9, 14.4, 18.0, 21.6, 29.7, 39.6, 50.4). e) Trial design of Experimental tasks illustrating perturbation of object rotation clockwise (red lines, CW) or counter-clockwise (blue lines, CCW) from initial position towards target angle by either 5.4°, 14.4° or 28.8°.

#### Grasping Studies – Calibration Task

At the start of each experiment, a calibration task was performed to estimate of the subject’s ‘switch-point’, defined by when there is a 50% chance of using either grasp. Due to individual differences in height and anatomical flexibility, the switch-point was variable between subjects; a determinate angle for one subject could be ambiguous for another. The switch-point therefore acted as an individualised ‘zero’ position for which the target angles of subsequent experiments were centred around. A total of 76 trials (randomised for each subject) across 19 unique target angles uniformly spanning 88.2° were used to estimate the switch-point (example calibration curve is shown in Figure 1c).

To initiate a trial, participants were asked to push down on the home-pad with their wrist positioned mid-pronation while lightly pinching their finger and thumb together. The length of time required to hold down the home-pad was uniformly varied from trial to trial (1 - 1.3 ms) to prevent participants acting on anticipation of the go cue. Upon hearing the go cue (auditory beep 500 Hz, lasting 50 ms), participants were asked to quickly reach out and comfortably grasp the target object between their thumb and index finger, pinching it lightly. After the precision grip, they returned to the home-pad to initiate the next trial sequence. If the home-pad was released too early (i.e. prior to the go cue), an error tone would sound (200 ms, 200 Hz) to indicate the subject should return to the start button.

For this task, participants wore LCD goggles (PLATO goggles; Translucent Technologies, Toronto, Canada) to occlude their vision until the object had rotated into place at the target angle. The goggles have a transition time from opaque to visible of 4 ms. Synchronously with the auditory go cue, the goggles transitioned from an opaque state to transparent to allow subjects to perform the prehension movement. In cases where participants wore visual correction glasses, the goggles were placed over the top and secured with a headband. Once the switch-point was established, the experimental tasks could commence.

#### Grasping Studies – Experimental Task

To investigate the influence of prior visual information on grasping in ambiguous conditions, the main task trial sequences were designed to first present subjects with the target object at a randomised initial angle for which they would not reach, but instead only observe. This was followed by the object rotating either 5.4°, 14.4°, or 28.8° clockwise (CW) or counter clockwise (CCW) to the target angle that they would reach for (Figure 1e), taking 10-50 ms depending on the shift magnitude. A total of 23 unique target angles spanning 50.4° either side of the measured switch point (Figure 1d) were presented in a randomised order, with each subject given their own randomisation. The rotation was concealed from subjects in the first, Static Experiment, and visible in Dynamic Experiments 1-3.

To reduce unnecessary sampling of the determinate region, target angles far from the switch-point (> ± 14.4°) were only given shift magnitudes of 5.4°. To further reduce experimental time and minimise repetition fatigue, the remaining targets were randomly given two of the three potential shift magnitudes, with a balanced distribution of all three magnitudes across the ambiguous range (≤ ± 14.4°). Finally, determinate angles were sampled at half the frequency of ambiguous orientations, with greater inter-angle spacing. All experiments consisted of a total of 216 trials.

Each experiment varied the timing of the go-cue relative to object rotation. A comparative timeline of each experiment’s sequence is shown in Figure 1b.

### Static Experiment

For the Static Experiment, participants were asked to wear LCD goggles with the purpose of blocking object rotation from the visual field to confirm the influence of previous object positioning on grasp choice.

Synchronous with subjects pressing the home-pad, the goggles transitioned to an opaque state before the object was rotated to the initial angle (Figure 1e). Once object rotation was complete, the goggles became transparent to allow subjects to view the initial angle for 1500 ms. After this delay, the goggles transitioned back to an opaque state and remained opaque for 650-950 ms, during which time the object rotated from the initial angle to the target angle. Once this was complete, the goggles converted to a transparent state (simultaneous with auditory go cue) to allow subjects to view and reach out to grasp the target angle.

### Dynamic Experiments

For all dynamic experiments, the goggles were removed to allow subjects to view object rotation from the initial angle to the target angle to explore how motion might change the influence of prior object position.

In these experiments, subjects viewed the initial angle for 1000-1300 ms before witnessing object rotation from the initial angle to the target angle. In Dynamic Exp. 1, the go cue sounded 50 ms prior to rotation onset. For Exp. 2, the go cue sounded 150 ms after rotation onset; this specific delay was chosen to allow time for visual processing delays (i.e. the time it takes for humans to perceive objects in their visual field (Hegdé, 2008)), allowing some, but limited, preparation before the go cue. For Exp. 3, this delay increased from 150 ms to 500 ms (Figure 1b). This delay was chosen to be large enough to allow subjects to fully prepare a movement in advance of the go cue, yet still brief enough for subjects to remain engaged with the task on each individual trial.

### Perception Study

Following the results of the grasping experiments, a second study was designed to investigate the perceptual consequences of observing a shift in object orientation, thereby investigating the mechanisms by which prior visual information may affect grasp choice. Here we wanted to test whether the same inverse bias we found in Dynamic Exp. 1-3 can be observed without the grasping action performed by the subjects, and thus can be attributed to perceptual, rather than motor planning, phenomena.

#### Participants

All participants were right-handed and naïve to each experiment and its predictions. In total, 11 healthy volunteers (20-22 years old, 7 females 4 males) were recruited for this study. There was no overlap in participation between this visual study and the previous grasping study, and all had normal or corrected to normal vision.

#### Perception Study - Task Sequence

In this study, participants were seated with set-up as described in *Grasping Studies – Instrumentation* and began with a calibration task in order to find an estimate of their centre of ambiguity. Following this, the main perception task was designed to investigate whether observing rapid object motion from an ‘initial’ position to ‘target’ leads to changes in how the target position is perceived by an individual. An overview of the trial sequence for this perception experiment is illustrated in Figure 4a.

As with the grasping task, participants initiated the trial sequence by holding down the home-pad. However, after observing object rotation from an initial angle to a target angle as in the Dynamic Experiments, participants were asked not to execute a grasp as before, but instead only continue to observe. After observing the target angle for 100 ms, vision was blocked by goggle opaqueness. This delay of 100 ms was chosen to allow sufficient time for perceptual processing (Hegdé, 2008), yet short enough that it is comparable to the time given to observe the target angle in the grasping experiments prior to the go cue. With subjects’ vision blocked, the object underwent another movement, concluding at a ‘test’ angle located either 5° CW/CCW (‘catch’ trials), or unchanged from the target angle orientation. To prevent participants from using auditory cues to distinguish between catch and non-catch trials, the object rotated on all trials before concluding at the final test angle position, however on only 10% of trials was the test position different from the target angle (i.e. a catch trial). This period of visual blocking lasted for 500 ms, chosen to ensure that any potential ‘after-image’ effects were absent. After this delay, vision was restored and participants were asked to report (by reaching to press keys on a keyboard placed in front of them) whether the object appeared to have shifted orientation between the target angle and test angle positions. If the test angle appeared CCW relative to the target angle, subject were asked to press ‘1’. If the test angle appeared unchanged from the target angle, ‘2’ was used. Finally, if it appeared CW, ‘3’ was pressed. Subjects were given unlimited time to observe the test angle and report their decision.

The prediction here being that if the inverse bias effect was due to perceptual phenomena, then subjects would be more likely to report that the object at the test position is further ‘CW’ relative to the target angle when the rotation from initial to target angle was CW versus CCW, despite the object not moving between the target and test positions. Here, no grasping action was required in order to allow investigation of solely perceptual phenomena.

### Data and Statistical Analysis

All analysis was performed in MATLAB using custom written scripts. For analysis of the contribution of the different experimental factors in the grasping task, a generalised linear mixed effect model (GLME) approach was adopted. GLME analysis allowed estimation of the contribution to grasp choice (C) of each experimental factor (target angle (TA), shift direction (Di), and shift magnitude (SM)), their pairwise interactions, as well as Subject (S) as a random effect. For reaction and movement time analysis, outliers were identified and excluded. Further detail is provided in Online Resource 1.

## Results

### Calibration Task

To begin each experiment, participants were asked to perform reach and precision grasp movements to a triangular prism presented at randomised target orientations. As expected, subjects transitioned from using Grasp 1 to Grasp 2 as the object sampled more clockwise angles (e.g. Figure 1c). Across all participants’ calibration sessions, the switch-point ranged between -58.1° and 32.8°, relative to the vertical axis, with varying degrees of sharpness (Halfwidth = 17.09 ± 2.15°); the calculation of halfwidth is provided in Online Resource 1 and exemplified in Figure 1c.

### Static Experiment

To confirm previous findings that prior information can bias grasp behaviour, the static experiment task added an additional component before grasping of the target angle: an initial angle which subjects would not reach for but only observe. Here, subjects were presented with this initial angle for a brief moment before LCD goggles occluded their view of the task, at which point the object rotated either CW or CCW to the target angle. Post-rotation, the LCD goggles then became transparent, triggering subjects to grasp the object at the target angle.

A visualisation of the generalised linear mixed effect model (GLME) fitted to the data for the static experiment is shown in Figure 2a-c. As expected from the design of the experiment, target angle was found to be a significant factor on grasp choice (CE = 2.734 (95% CI [2.389, 3.081]), t-Stat = 15.5, dF = 2150, p < 0.001). Furthermore, there was a significant effect of shift direction on grasp choice (CE = 0.417 (95% CI [0.183, 0.652]), t-Stat = 3.5, p < 0.001), suggesting that the orientation of initial angle influenced the planning of the grasp for the target angle, i.e., the presentation of the object at an initial position that would likely require Grasp 1 increased the likelihood of subjects then later adopting Grasp 1 for the ambiguous target angle. The sizes of both these significant effects are above the minimally detectable effect with a power of 0.8. To gain an intuitive understanding of how grasp choice is different between CW and CCW object rotation directions, we calculated from the model prediction the difference in probability of Grasp 2 at the switch point angle (indicated by black lines on Figure 2a-c). For shift magnitudes of 5.4°, it was 0.08, whereas for larger shift magnitudes of 14.4° and 28.8°, the difference was larger at 0.11 and 0.15. However, neither shift magnitude nor any interactions were found to be significant (SM CE = -0.030 (95% CI [-0.186, 0.125]), t-Stat = -0.4, p = 0.702; TA*Di CE = 0.101 (95% CI [-0.397, 0.599]), t-Stat = 0.4, p = 0.691; TA*SM CE = 0.137 (95% CI [-0.124, 0.398]), t-Stat = 1.0, p = 0.304; Di*SM CE = 0.112 (95% CI ([-0.112, 0.336]), t-Stat = 1.0, p = 0.325).

**Figure 2.**
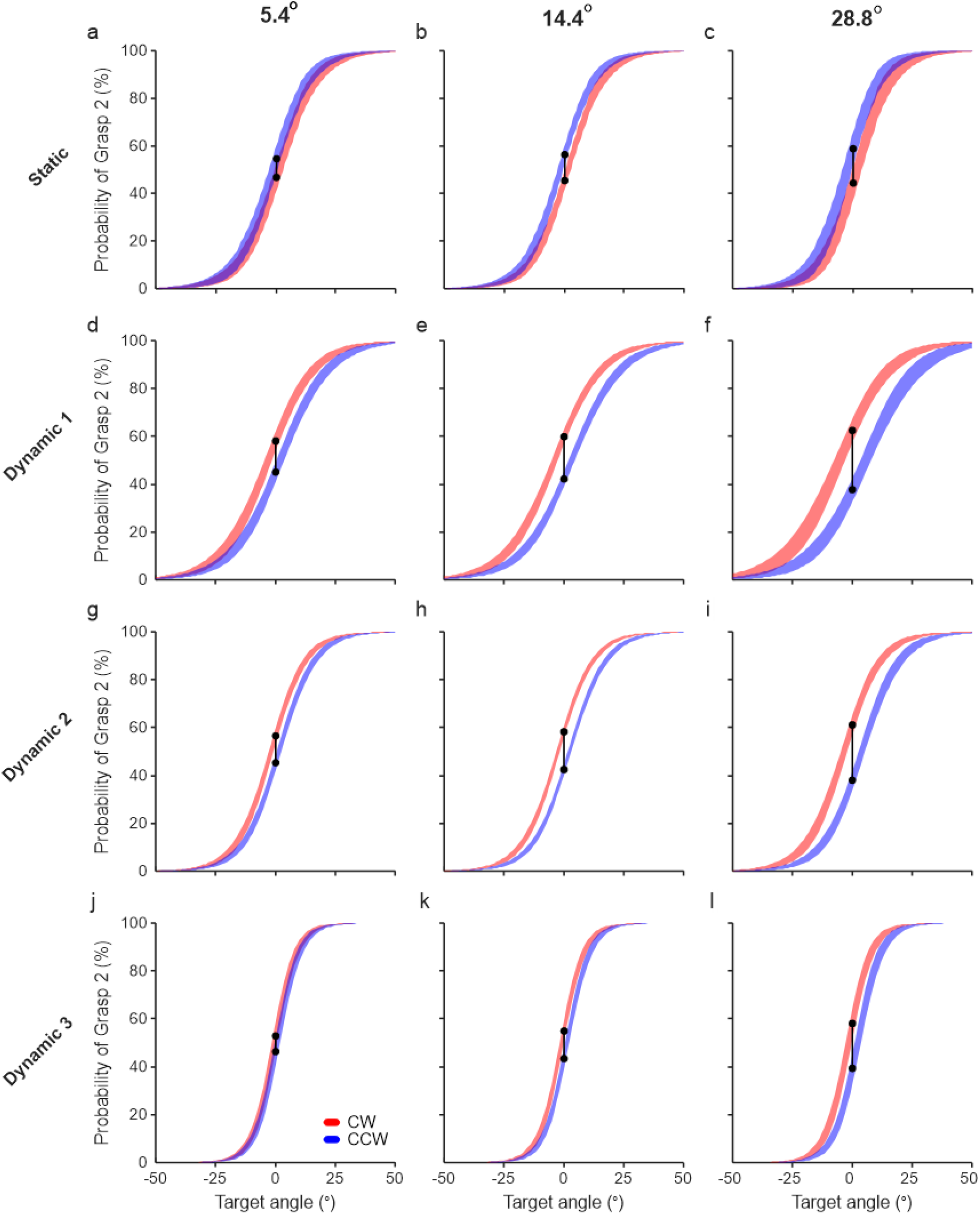
GLME visualisation from all subjects of the Static Experiment (a-c), Dynamic Exp.1 (d-f), Dynamic Exp. 2 (g-i) and Dynamic Exp. 3(j-l). Choice is predicted separately for each rotation direction (red and blue lines) and shift magnitude (5.4° shown in (a,d,g,j), 14.4° shown in (b,e,h,k), 28.8° shown in (c,f,i,l)) for target angles 50° either side of the centre of ambiguity. a-c) GLME model for the static experiment: C ∼ TA + Di + SM + TA:Di + TA:SM + Di:SM + (1|S). d-l) GLME model for combined dynamic experiments 1-3: C ∼ TA + Di + SM + DE + TA:Di + TA:SM + TA:DE + Di:SM + Di:DE + SM:DE + (1|S). The SEM across all subjects is illustrated by line thickness. The difference in probability at 0° for each rotation direction is illustrated with vertical black lines at 0° target angle. C, choice; TA, target angle; Di, direction; SM, shift magnitude; DE, delay duration; S, subject; CW, clockwise; CCW, counter-clockwise.

To summarise, the influence of prior object position on grasp choice at an ambiguous target is in agreement with the priming effects previously reported in the literature (Gallivan et al., 2015, Gallivan et al., 2016). Furthermore, there is weak evidence to suggest that the higher the likelihood of the initial position requiring Grasp 1, the stronger the effect of subjects then later adopting Grasp 1 for the ambiguous target angle, however this was not statistically significant.

### Dynamic Experiment 1

To investigate any possible influences that prior dynamic visual information could have on grasping in ambiguous settings, the following experiment kept the same conditions as the static experiment, but without the LCD goggles, allowing subjects to watch the object rotate from the initial angle to the target angle.

As expected, the target angle was found to be a significant factor (CE = 2.156 (95% CI [1.902, 2.410]), t-Stat = 16.6, dF = 2554, p < 0.001). The direction of object rotation was also found to be a significant factor (CE = -0.699 (95% CI [-0.910,-0.488]), t-Stat = -6.50, p < 0.001), yet the effect was opposite to that observed in the Static Experiment, i.e., the presentation of the object at an initial position that would likely require Grasp 1 increased the likelihood of subjects then later adopting Grasp 2 for the ambiguous target angle, referred to hereon as the ‘inverse bias effect’. The sizes of both these significant effects are above the minimally detectable effect with a power of 0.8. To gain an intuitive understanding of how grasp choice is different between CW and CCW object rotation directions, we calculated from the model prediction the difference in probability of Grasp 2 at the switch point angle (indicated by black lines on Figure 2d-f). For shift magnitudes of 5.4°, it was -0.14, whereas for larger shift magnitudes of 14.4° and 28.8°, the absolute difference was even greater at -0.17, and - 0.23, respectively. However, neither shift magnitude any interactions were found to be significant (SM CE = 0.066 (95% CI [-0.078, 0.211]), t-Stat = 0.9, p = 0.369; TA*SM CE = -0.137 (95% CI [-0.314, 0.039]), t-Stat = -1.5, p = 0.126; Di*SM CE = -0.172 (95% CI [-0.373, 0.029]), t-Stat = -1.7, p = 0.094).

To summarise these findings, a clockwise rotation to an ambiguous target angle increased the probability of Grasp 2 (a grasp non-congruent with the initial position) at the switch-point in comparison to Grasp 1 (a grasp congruent with the initial rotation), while a counter-clockwise motion increased the likelihood of Grasp 1. Although the difference between grasp choices appears larger for larger shift magnitudes (Fig. 2d-f), both effect sizes for rotation magnitude and its interaction with direction were smaller than the estimated minimal effect size at the power of 0.8, given the sample size. This is a potential reason for both factors being non-significant.

### Dynamic Experiments 2 and 3

To investigate whether the timing of the go-cue relative to object rotation can influence the strength of the inverse bias effect, this delay was varied from -50 ms in Dynamic Exp. 1, to 150 and 500 ms in Exp. 2 and 3 respectively (see Figure 1b). We predicted that the longer subjects have to observe the target angle before receiving the go cue, the inverse bias effect would become weaker, owing to the prior information becoming more historical and therefore less influential. Thus, data from Exp. 1, Exp. 2 and Exp. 3 were combined into one model with delay was added as a factor into the GLME.

The visualisation of the results of the model fitted to the data when the go cue was delayed until 150 ms and 500 ms post-rotation are shown in Figure 2g-i and 2j-l respectively, with the results for a delay of -50 ms shown in Figure 2d-f. Target angle again remained significant (CE = 3.224 (95% CI [3.007, 3.442]), t-Stat = 29.0, dF = 6840, p < 0.001), as expected. The direction factor also remained significant (CE = -0.582 (95% CI [-0.721, -0.443]), t-Stat = -8.2, p < 0.001). There was no significant effect of go cue delay on choice (CE = -0.077 (95% CI [-0.177, 0.022]), t-Stat = -1.5, p = 0.128), nor a significant direction-delay interaction (CE = 0.104 (95% CI [-0.036, 0.243]), t-Stat = 1.5, p = 0.144).

Though this latter observation is suggestive of a constant inverse bias effect for all planning times, we cannot rule out the possibility of a small effect that was not significant due to smaller sample size. Indeed visually, the difference in grasp choice probability at the switch point between CW and CCW directions narrows, decreasing from 0.13 to 0.07 for shift magnitudes of 5.4°, from 0.17 to 0.12 for 14.4° shifts, and 0.25 to 0.19 for 28.8° shifts (Figure 2d-l). Importantly, there was a significant interaction between target angle and delay (CE = 1.00 (95% CI [0.826, 1.182]), t-Stat = 11.1, p < 0.001), indicating a steeper transition between Grasp 1 and Grasp 2, suggestive of greater certainty in grasp choice with greater planning time. The sizes of all significant effects described here are above the minimally detectable effect with a power of 0.8. To quantify this, we estimated the change in Halfwidths from the individual subjects’ GLME predictions. Between delays of -50 ms, 150 ms and 500 ms, the Halfwidth reduced from 20.8° to 15.8° to 10.2° for CW and from 21.6° to 16.3° to 10.3° for CCW. Shift magnitude did not have a significant effect on grasp choice, however, a significant interaction between direction and shift was found (CE = -0.199 (95% CI [-0.331, -0.067]), t-Stat=3.0, p = 0.003), indicative of a stronger bias effect with a larger rotation between the initial and target angles. None of the other interactions were significant (TA*Di CE = -0.086 (95% CI [-0.359, 0.187]), t-Stat = -0.6, p = 0.535; TA*SM; CE = -0.078 (95% CI [-0.213, 0.057]), t-Stat = -1.1, p = 0.257; DE*SM CE = 0.003 (95% CI [-0.064, 0.069]), t-Stat = 0.1, p = 0.941).

To summarise, when the delay between target angle presentation and the go cue increased from -50 to 500 ms, subjects showed less ambiguity in their grasp choice. Furthermore, the inverse bias effect persisted even with greater planning time allowance and was influenced by the magnitude of rotation.

### Reaction and Movement Times

When completing each trial, participants were instructed to act quickly but with priority to accuracy and comfort. As such, each experiment was not designed as a RT task. Yet, one might predict that the greater the uncertainty induced by the experimental conditions, the longer the RT (Wood and Goodale, 2011). Indeed, significant differences in RTs were observed both within (ambiguous versus determinate target angles) and across (go cue timing) experiments.

To quantify the relationship between each of our behavioural measures and delay timings, three separate GLME analyses were performed with delay as the sole (continuous) variable and estimated values of HW, mean reaction and movement times as each independent variable (Figure 3a-c). Each analyses showed a significant effect of delay on halfwidth (DE CE = -0.406 (95% CI [-0.652, -0.161]), t-Stat = -3.4, dF = 30, p = 0.002), reaction time (DE CE = -0.476 (95% CI [-0.727, -0.225]), t-Stat = -3.9, dF = 30, p < 0.001), and movement time (DE CE = -0.248 (95% CI [-0.438, -0.059]), t-Stat = -2.7, dF = 30, p = 0.012). The effect sizes described here are close but below the minimally detectable effect with a power of 0.8, yet are significant (p < 0.05). Given the sample size, this might be a false positive; however, it might also be a genuine effect present in our data.

**Figure 3.**
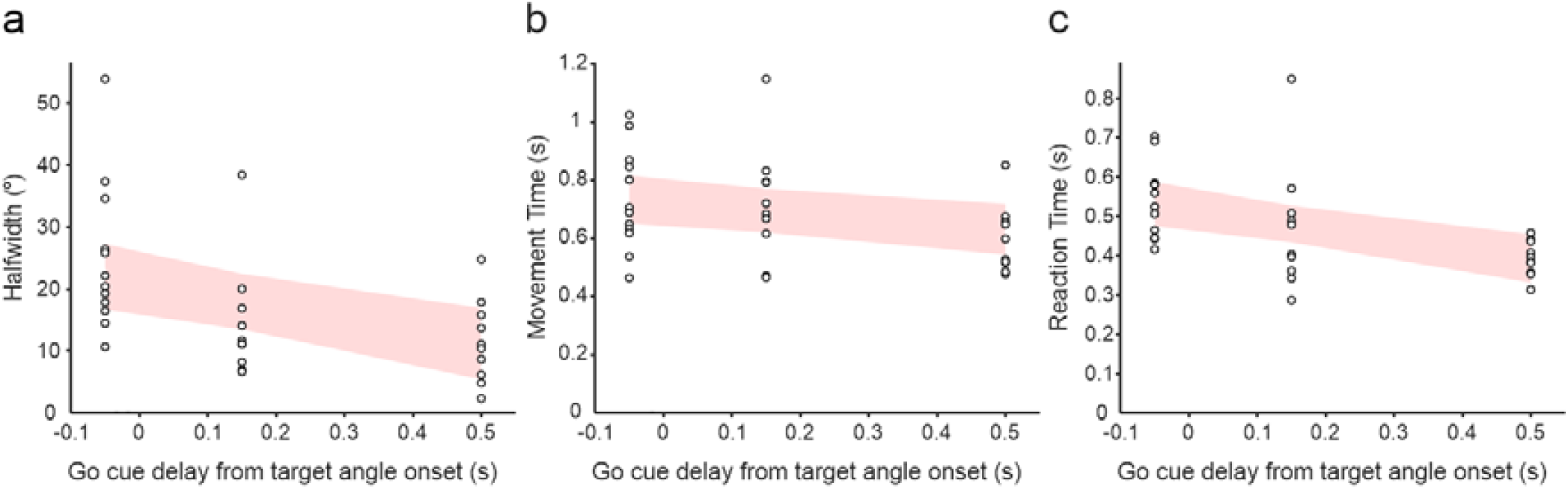
Effect of delay from target angle onset to go cue on behavioural measures. a) Halfwidth was estimated for each subject and delay, predicted from individual GLM analysis (GLM model: C ∼ TA). GLME model: HW ∼ 1 + delay + (1|S). b) Mean movement times were calculated following outlier removal for each subject and delay. GLME model: MT ∼ 1 + delay + (1|S). c) Mean reaction times were calculated following outlier removal for each subject and delay. GLME model: RT ∼ 1 + delay + (1|S). Shading illustrates prediction of each model with confidence intervals at 95%.

To investigate the effect of target angle on RTs, a LME approach was used. In agreement with previous work, RTs for target orientations closer to the centre of ambiguity were greater than those more determinate (log transformed raw values are shown in blue in Online Resource 2). To summarise these experiments, as the experimental delay increases, there is both greater uncertainty as shown by the halfwidth measure, as well as slower the RTs within and across experiments.

### Perception Task

Given the finding that the directionality of object rotation can influence grasp choice, we investigated whether this could be attributed to changes in how the target position is perceived after observing object rotation. In this secondary study, subjects were asked to observe object rotation from initial to target angles as for the Dynamic Experiments, except after target angle presentation, visual input was briefly blocked. Subjects were then re-shown the object at a test angle and asked to report whether it appeared further CW, CCW, or ‘unchanged’ from the target angle position (Figure 4a). On non-catch trials, the test angle was identical to the target angle, and thus a ‘correct’ answer would therefore be ‘Unchanged’.

**Figure 4.**
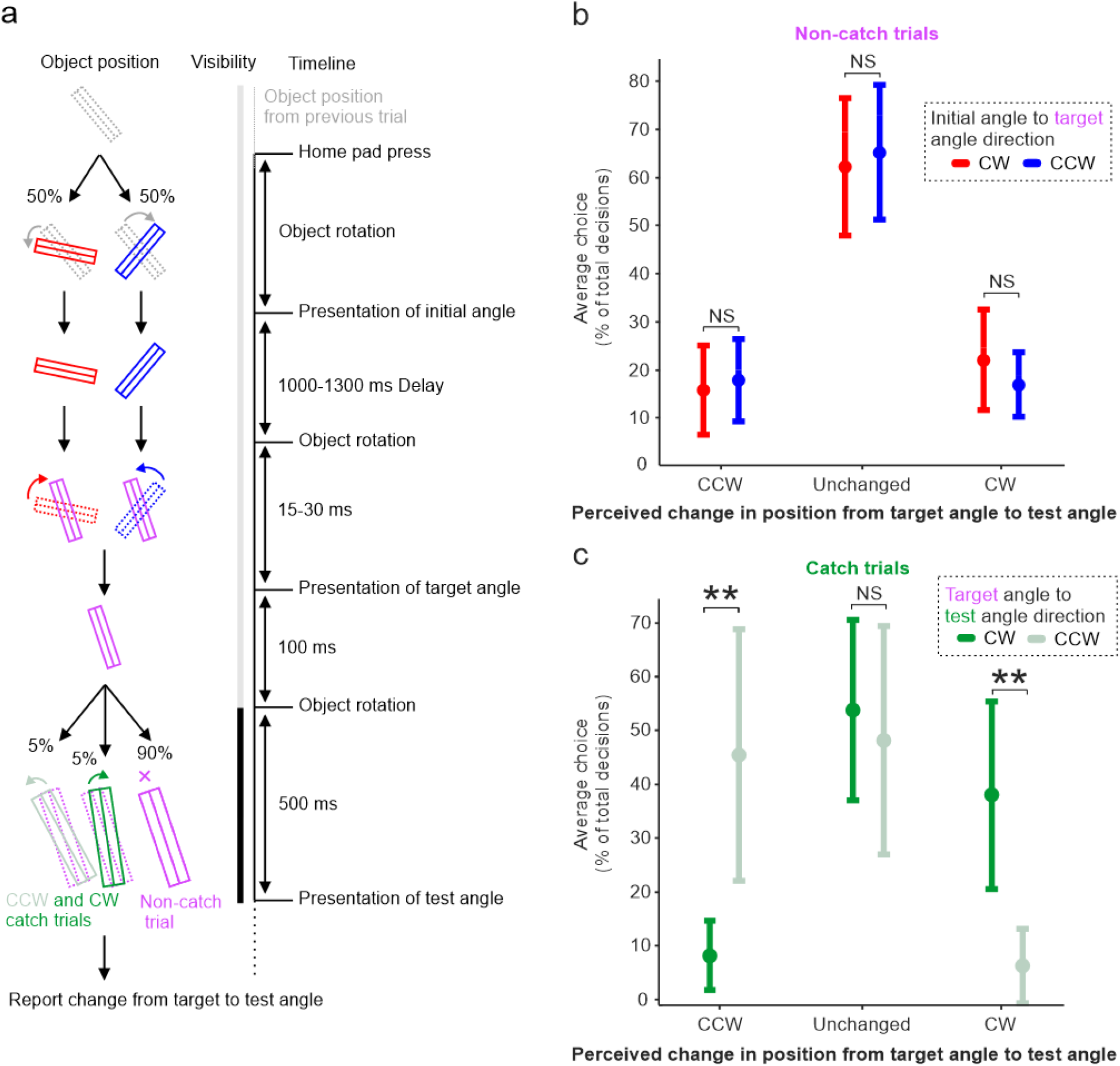
Experimental timeline and behavioural results from the Perception Experiment. a) Timeline of a single trial during the perception experiment. Periods of object visibility and obscurity are illustrated by light and dark bars respectively. The time given to observe the target angle was 500 ms, whilst the duration of visual blocking before test angle presentation was 1 s. b) Following observation of object rotation (CW or CCW) from an initial angle to target angle, then subsequent presentation of the test angle, a decision of ‘CCW’ reflects that the subject perceived the test position to be further counter-clockwise than the target position, whilst ‘CW’ reflects that it appeared further clockwise. Here, averaged perceptual decisions are taken only from trials wherein the test position was unchanged from the target position. c) Averaged perceptual decisions taken only from trials wherein the test position was different from the target position, following either CW or CCW rotation from the target angle to test angle. The error bars represent confidence intervals of 95%. NS: No significant difference. ** p < 0.01.

Average choices on non-catch trials (i.e. target angle = test angle) are shown in Figure 4b. There was no significant difference between the error rate in reporting ‘CCW’ (p = 0.296) when the rotation direction from initial to target angle was CW versus CCW, nor in reporting ‘CCW’ (p = 0.253). *Thus, there does not seem to be a bias in perceived object position at the target angle following object motion from the initial angle. This is consistent with a non-perceptual explanation for the inverse bias effect observed in the grasping experiments, however potential contributions cannot be fully ruled out.*

To confirm that participants were attending to and comparing test angle position relative to target angle, catch trials were included (Figure 4c). Here, the object, rather than remaining at the target angle following the initial angle to target angle transition as for the above, rotated to the test angle.

When this rotation was CW, subjects were on average significantly more likely to correctly report that the test angle appeared CW relative to the target position than CCW (CW 38.01% versus CCW 6.36%, p = 0.0056), and significantly more likely to correctly report CCW than CW on a CCW rotation (CW 8.26% versus CCW 45.45%, p = 0.0054). The error rate on catch trials was 53.72% for CW shifts and 48.18% for CCW, a reflection of a task difficulty that was necessary to induce in order to investigate perceptual phenomena on non-catch trials. These findings confirmed that subjects were attending to the task, and that the lack of perceptual bias found on non-catch trials cannot be attributed to lack of engagement.

## Discussion

In this study, subjects were placed into ambiguous settings with an object that afforded one of two precision grasps (Grasp 1 or Grasp 2) in order to compare the influence of static and dynamic visuomotor priming on motor ambiguity. In alignment with previous priming studies (Seegelke et al., 2016, Borghi et al., 2007, Vainio et al., 2007), we demonstrate that the presentation of an initial, priming orientation of the object (at either a determinate or ambiguous orientation) was sufficient to influence choice of grasp when the object was then shown at an ambiguous orientation (Static Experiment). More specifically, subjects’ grasp choices for the target angle were biased toward the orientation congruent with what they would have likely used for the initial position. However, when rotation of the object from the initial angle to the target position was visible, a significant but inverse effect of rotation directional bias on grasp choice was found (Dynamic Exp. 1-3). Here, a clockwise rotation from an observed Grasp 1 position to an ambiguous angle increased the likelihood of Grasp 2, and vice-versa with a counter-clockwise rotation, referred to here as the ‘inverse bias’ effect (Prediction 3).

One of our original predictions stated that object motion could prevent proactive influence (Prediction 2), i.e. motor planning would begin only at time of target angle presentation and there would be no effect of initial position on the grasp chosen, explained by the ‘dorsal amnesia hypothesis’. However, in line with previous findings (for review see (Schenk and Hesse, 2018)), we find no evidence in support of this view in our experimental context.

Upon establishment that visible rotation and its directionality influenced grasp choice, two further experiments were designed to explore its persistency over time. Increasing the delay between rotation onset and the go cue to both 150 ms and 500 ms still produced the inverse bias effect, and although the delay duration was not a significant contributor to the strength of the effect, the separation of grasp choice probability between CW and CCW rotation directions did narrow with greater planning time. This indicated a non-significant reduction in the strength of the inverse bias effect with greater planning time, however, this observation may reflect the constraints of the experimental design, namely the evaluation of only three delay intervals in separate subject groups.

Despite not enforcing subjects to react as quickly as possible (accuracy and comfort of the grasp were emphasised during task instructions), the RT still reflects the time needed for the cortical circuitry to choose between grasps (Wood and Goodale, 2011, Rosenbaum et al., 1992). RTs decreased significantly with increasing delay (Figure 3c), suggesting that preparatory processes were engaged during the delay interval.

In addition to the inverse bias effect, another feature of observing object motion was its influence on grasp choice uncertainty. This was revealed by a significant interaction between target angle and delay, as well as the direct comparison of halfwidth decreasing with greater delays (Figure 3a). Visually, this appears as a shallower transition between the two grasp choices the earlier the go cue was given, a consequence of increased variability suggestive of more inefficient planning (Dijkerman et al., 2009). When allowed greater planning time post object motion, uncertainty reduces and the inverse bias effect potentially weakens.

The mechanisms by which object motion induces these signs of increased uncertainty, and how they may be linked to the inverse bias effect require further investigation to uncover. We suggest that given for Dynamic Exp. 1 the go cue was triggered immediately before object rotation onset, the initiation of movement planning prior to knowing the target orientation could explain the greater uncertainty and lead participants to incorrectly predict the final target orientation.

Predicting the future location of a moving object is a familiar task, yet some consistent biases have been reported. It could be considered a good strategy for example, that if an occluded object is approaching an observer, to overestimate its speed in order to be cautious and avoid potential collision (Schiff and Oldak, 1990). Contrastingly, when asked to intercept an occluded object moving parallel to the observer, its speed is often underestimated, causing people to intercept its path too late (e.g. (Sokolov and Pavlova, 2003, Peterken et al., 1991, Battaglini and Ghiani, 2021)). An overestimation of the final target position would push subjects towards planning a grasp different from the grasp that would have been used for the actual target position, and therefore could be a mechanism for the inverse bias effect. In Dynamic Exp. 2 and 3 (wherein the go cue is given after rotation is finished), prediction of the final object orientation is arguably unnecessary and could be planned anew when the go cue is given as per the dorsal amnesia hypothesis, yet this does not seem to be true. Instead, the inverse bias effect persists, suggesting that the plan to grasp is not updated at the go cue, but rather remains similar to the one created during object rotation.

An alternative suggestion for why grasp selection in ambiguous settings is sensitive to target rotation could be that rapid object rotation leads to momentary changes in how the target angle is visually perceived. Visual perceptual phenomena are not unfamiliar to the human mind, for example, the Fröhlich effect and representational momentum both describe visual illusions in which the location of moving objects are misperceived (Müsseler et al., 2002, Kanai et al., 2004, Fu et al., 2001, Merz et al., 2023, Hubbard, 2018). In our case, if the target orientation is perceived to have rotated further than is true, the bias effect could then be attributed to a perceptual error that degrades with greater time given to observe the target position prior to grasping. If a perceptual error is present, we hypothesised that, participants’ perception of final object position would be affected by rotation direction from the initial angle to target. Yet, the proportion of error decisions of ‘CCW’ compared to ‘CW’ was not influenced by rotation direction. This is consistent with a non-perceptual phenomena explanation for our inverse bias effect, and may instead suggest that the bias effect is linked to motor planning, though, the contribution of other perceptual phenomena cannot be ruled out and requires further investigation.

It has been shown that in situations affording more than one possible action, the brain prepares multiple competing movements before selecting one (Stewart et al., 2013, Cisek, 2007). Indeed, when presented with multiple reach targets and forced to act before knowing the final location, the initial movement vector reflects an average of the competing motor plans (Stewart et al., 2014, Haith et al., 2015). It has also been demonstrated that sequential presentation of potential targets can lead to proactive influence, i.e. the movement preparation associated with the first target can influence the motor plan for a secondary target (Gallivan et al., 2016). In Gallivan *et al.*, 2016, (Experiment 2) participants were cued to grasp one of two potential targets after they had each been sequentially presented. This led to subjects becoming biased to adopt a grasp for the cued ambiguous target that was congruent with a determinate angle shown prior to (proactive co-optimisation) or after (retroactive co-optimisation) the ambiguous target. The bias effect observed with our Static Experiment is consistent with this proactive co-optimisation, whereby the choosing of Grasp 2 is more likely if the object was rotated from an initial position greater in congruency with Grasp 2, and likewise for Grasp 1. It is curious, however, that for all our experiments, the trials that began and ended ambiguous still produced a bias effect. Though Gallivan and colleagues showed that in their study a similar trial type yielded no bias, a key experimental difference is that whilst they showed the exact same ambiguous orientation, ours changed by 5.4°. Thus, one interpretation of this could be that contrast in object orientation between the initial and target position, or in other words contrast between differing choice certainties, is needed to induce this bias effect.

Regarding the inverse bias effect observed in Dynamic Exp. 1-3, some previous studies have reported inverse-like effects in response to priming stimuli. In one study, participants enlarged their finger aperture after the presentation of a small object when reaching to grasp a successive medium sized object, and shrank their finger aperture after observing a large prime object (Pisu et al., 2020), an outcome that is likely a consequence of the Uznadze illusion (Uccelli and Bruno, 2024, Uccelli et al., 2019). Furthermore, this effect on grip aperture persists when participants must grasp without vision, yet effect strength decreases rapidly throughout the movement when the object remains in full-vision (Uccelli and Bruno, 2024). This is consistent with a qualitative observation that the inverse bias effect is reduced for all shift magnitudes with longer preparation times (Figure 2d, g, i), though a modified experimental design is needed to establish the statistical significance of this finding. Still, our inverse bias effect could be interpreted as evidence of an Uznadze-like aftereffect in an ambiguous grasp setting.

If indeed two neural populations within the grasping network are competing for each grasp (i.e. each are planned simultaneously) in situations of motor ambiguity, perhaps the observation of the object moving away from the preferred position of one population first inhibits its activity, before facilitating activity corresponding to the alternate grasp. The antagonism of potential actions is known to be subject to bias from a variety of both cortical and subcortical inputs, with the competition playing out across a reciprocally interconnected fronto-parietal system (Cisek and Kalaska, 2005, Matelli and Luppino, 2001, Borra et al., 2017). Evidence favouring a given choice causes activity to increase for its neural population, while information against a choice instead causes suppression of its neural population (Cisek, 2007). An increase in available information would under normal circumstances reduce RTs (Smith and Ratcliff, 2004, Ratcliff et al., 2003), and indeed, in our study, we observed a slowing in RTs with increasing delay. That being said, the quality of evidence can influence changes in RTs, and so adding available information (observation of the object rotation) may not enhance processing but rather complicate a decision by forcing further processing across the action selection network, or by introducing noise that outweighs any additional signal gained from un-occluded vision, hence leading to greater RTs at a delay of -50 ms (Ratcliff and McKoon, 2008, Palmer et al., 2005). By processing, we mean here disruption of the initial plan to grasp the object congruently with the initial orientation, in favour of planning a grasp congruent with predicted end object position – a consequence of observing object motion.

In conclusion, whilst static visuomotor priming has been shown to facilitate the convergence of action plans into one singular intent facilitating motor execution, the mechanisms of grasp selection in ambiguous settings can also be influenced, and potentially disrupted, by observation of object motion prior to grasp execution. This effect may not be due to certain perceptual phenomena, but could potentially be attributed to a consequence of cortical inhibitory circuits involving competing neural populations. Further experiments aimed at disrupting motor planning processes might help delineate the neural mechanisms behind this inverse bias. Understanding these neural mechanisms, and in particular initial inhibition underlying this motor bias could have implications for neurodegenerative diseases that exhibit motor planning and execution deficits, for example Parkinson’s Disease (Maffitt et al., 2025) and Dystonia.

## Acknowledgements

The authors thank Mr Norman Charlton for mechanical engineering of the experimental setup, and Prof. Stuart Baker for designing the contact circuit and initial Sequencer scripts. For the collection of pilot data, the authors would like to thank Ms Annabelle Clark and Ms Amn Ashhad. We would also like to thank Dr Steven Jerjian for critical feedback on earlier versions of the manuscript and Prof. James Kilner for suggestions to study design.

