## Supplementary Methods for "From static to dynamic: how object rotation influences grasp decisions in ambiguous settings"

Experimental Brain Research

Corresponding Author

Dr Alexander Kraskov, Biosciences Institute, Newcastle University, Newcastle Upon Tyne, UK, NE2 4HH,

### **Online Resource 1 - Supplementary Methods**

### **Data and Statistical Analyses**

Grasp choice data were gathered using the custom designed contact circuit and sampled at 5 kHz (CED Micro 1401, Cambridge Electronic Design) using Spike2 software. All data analyses and statistics were performed using custom written MATLAB scripts. For the grasping task, reaction time (RT) was measured as time from go cue to home pad release. Movement time (MT) was measured as time from release of home pad to contact with the target object.

#### **Calibration Task**

Choice of grasp was coded with reference to which edge of the target object the thumb made contact with. For the calibration experiments, the probability of using a ‘Grasp 2’ was calculated for each target angle and a logistic regression curve was fitted to the data using the function below, where *x_o_* is the subject’s switch point and *k* is a scaling parameter.

$$f\left( x \right)=\frac{1}{1+e^{-k(x-x_{o})}}$$

This allowed two metrics to be calculated, the switch point and halfwidth. The angle for which there is a 50% chance of using either grasp was used as an estimation of each participant’s switch point (=*x*_o_), later referred to as the 0° angle (Figure 1b). The halfwidth provided a measure of switch sharpness, calculated as the difference between target angles for the 75% and 25% choice probabilities (=*2ln(3)/k*). This was used as an indicator of subject uncertainty across the task as a whole.

#### **Grasping Experimental Tasks**

For analysis of the contribution of the different experimental factors, a generalised linear mixed effect model (GLME) approach was adopted. We selected a GLME approach because it allows the modelling of group level variation, suitable for repeated measures and supports random effects. Logistic regression was the natural choice since our outcome measure is binary (grasp1 vs grasp2), and the experiment was set up in such way (i.e. choice of midpoint during calibration procedure) that a clear sigmoid was observed in behavioural (choice) data. GLME analysis allowed estimation of the contribution to grasp choice (C) of each experimental factor (target angle (TA), shift direction (Di), and shift magnitude (SM)) their pairwise interactions and triple interaction. The factors and interactions included into the model are based on our predictions. We expect a strong effect of the target angle due to the design of the experiment, wherein target angles were selected around the switch point for each individual subject. Target angles for each subject were shifted to zero midpoint, estimated by fitting GLM to individual subject data with target angle as the only predictor. We checked for absence of over and under dispersion (dispersion was 1 for static and dynamic models), we also checked inflated variance in the data (VIF); each variable had a VIF value less than 3.

For the Static Experiment, we expected a priming effect similar to previous literature, in our case in the form of shift direction (Garrido-Vásquez et al., 2021, Hesse et al., 2008, Roche et al., 2015, Gallivan et al., 2015, Gallivan et al., 2016). For shift magnitude, we expected a larger effect for greater changes in orientation, i.e. the more determinate the initial orientation, the greater the influence on grasp choice for the target angle. We also predicted that behavioural bias effects would be stronger for the ambiguous range of orientations than for determinate orientations, therefore interactions TA:Di, TA:SM were included into the model. For the inclusion of the Di:SM interaction, we hypothesised that larger shifts would have a greater impact on the effect of object rotation direction. Subject specific variation (S) was accounted for by its introduction as a random intercept effect into the model below.

C ~ TA + Di + SM + TA:Di + TA:SM + Di:SM  + (1|S)

To give an indicator of the effect size that can be detected given a significance level of 0.05 and a fixed sample size of 10 subjects, we ran a simulation referred to as effect-size sensitivity (Giner-Sorolla et al., 2024). At a power of 0.8, the minimal detectable effect size for each GLME factor for the static experiment was at least 0.3 (TA = 0.27, Di = 0.27, SM = 0.2, TA:Di = 0.27, TA:SM =0.27, Di:SM = 0.29). An effect size of 0.3 is typically interpreted as a small effect (Chen et al., 2010), meaning that our study should be sufficiently powered to detect at minimum small effects.

For Dynamic Exp. 1, the model was constructed the same as for the Static Experiment, with the only difference being the prediction behind the factor of direction.

For Dynamic Exp. 2 and 3, we hypothesised that the greater the delay between the end of object rotation, i.e. revelation of the target angle, and the go cue, the less uncertainty there would be in the grasp configuration choice. We further hypothesised that a greater delay would result in a weakened effect of direction on grasp choice, i.e. the more historic the prior information, the less influential it becomes. To investigate this, an additional factor of delay time (DE) was introduced into the model, coded as -0.05, 0.15, and 500 ms for Dynamic Exp. 1, 2 and 3 respectively.

C ~ TA + Di + SM + DE + TA:Di + TA:SM + Di:SM + TA:DE + Di:DE + SM:DE + (1|S)

The minimal detectable effect size using sample sizes of 12, 10, and 10 for each dynamic experiment with alpha = 0.05 and a power of 0.8 was at least 0.19 for each GLME factor, thus this study should be sufficiently powered to detect at minimum small effects.

For each experiment, the model was fitted in MATLAB using fitglme function (MathWorks, 2024). The target angle variable was shifted to the zero midpoint estimated from fitting a simple GLM to each subject individually (C ~ TA). Direction was coded as a categorical variable (CW, 0; CCW, 1) whilst target angle and shift magnitude were numerical. The shift magnitude and delay variables were mean-centred. All continuous variables were z-scored for easier interpretation of beta coefficients. A GLME with binomial distribution and logit link function was used. Model parameters were estimated using maximum likelihood with approximate Laplace method. T-statistic values (t-Stat), coefficient estimates (CE) and confidence intervals (CI) were obtained for each factor. The threshold of T-values were set to a p-value of 0.05 for each experiment. Statistics throughout are given as mean ± SE. For easier visualisation of GLMEs, target angles were restored from z-scoring.

#### **Grasping Task: Reaction and Movement Time Analysis**

Previous work has shown that both RTs and MTs are longer for ambiguous grasps than determinate (Wood and Goodale, 2011). To test for this in our grasping experiments, we used an LME approach to explain our RT and MT data using the absolute difference of the target angle from the ambiguity point as the single factor in the models below:

RT Model: ln RT ~ |TA-X_o_| + (1|S)

MT Model: ln MT ~ |TA-X_o_| + (1|S),

where X_o_ is a switch point estimated by fitting a sigmoid function for each subject separately. A significant linear decrease in these models would correspond to longer RT/MT for ambiguous target angles in comparison to determinate angles. Reported effects are estimated from each model as the difference between the centre of ambiguity, 0°, and ± 40° from this switch-point, and then reversed transformed from log values. A cut-off of 40° was used due to the decreased sampling rate observed at extreme angles from the centre of ambiguity. Trials for which RT was longer than 1.5 s or shorter than 150 ms were excluded from analysis; MTs however had no explicit high/low cut-off. For both ln RT and ln MT analysis any remaining trials outside the mean ± 3 SD were also excluded - this was then repeated recursively until no more trials could be excluded. On average across all 4 experiments, 2.6% of the trials were discarded for reaction time analysis and 1.3% for movement time analysis.

#### **Perception Experimental Tasks**

Perceptual data were obtained from subjects’ key presses of ‘1’ (test angle appeared CCW to target angle), ’2’ (appeared unchanged), or ’3’ (test angle appeared CW to target angle).

Trials for which the test angles were indeed shifted CW or CCW from the target angles acted as catch trials, with the remainder trials (i.e. test angle = target angle) denoted ‘unchanged’. The catch trials were introduced to make sure the subjects do pay attention to the task and detect correct shifts when these shifts were indeed present.

For unchanged trials, the results were first split by directionality of object rotation from initial angle to target angle, dividing the data into CW and CCW trials. Given the frequency of trials for each target angle varied according to level of ambiguity, matching the grasping task, the subject decisions (i.e. CCW, Unchanged, CW) per test angle were expressed as a percentage. The percentage of each decision was then averaged across all test angles. To investigate the effect of rotation directionality on object position perception, paired t-tests were used to compare CW and CCW conditions for each decision.
