## Supplementary Figures for "From static to dynamic: how object rotation influences grasp decisions in ambiguous settings"

Experimental Brain Research

Corresponding Author

Dr Alexander Kraskov, Biosciences Institute, Newcastle University, Newcastle Upon Tyne, UK, NE2 4HH,

### **Online Resource 2 - Supplementary Figures**

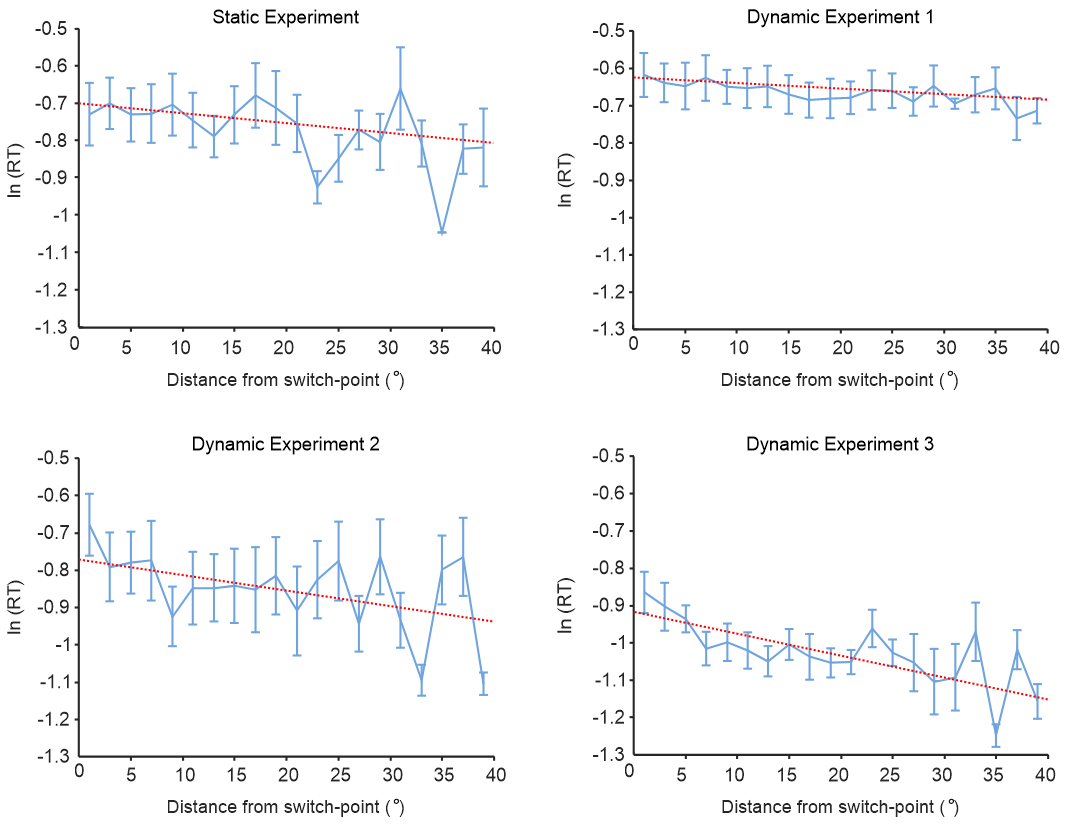

***Online Resource 2. Reaction times within both Static Experiment and Dynamic Exp. 1-3****. Reaction times for each target angle were measured from time of go cue to release of home-pad and averaged as angular distance from the switch point and across subjects. Log transformed values of the averages are shown in blue (ambiguous angles are emboldened). To illustrate the effect of distance from the switch point on reaction time, GLME analysis was carried out for each experiment; estimated values for each distance are shown by the red lines. The LME-estimated decrease in RTs from the switch point (0°) to 40° was 30.9 ms for Dynamic Exp. 1, 70.5 ms for Exp. 2, 83.7 ms for Exp. 3, and 50.0 ms for the Static Experiment. Target angle had a significant effect on reaction time for all experiments (Dynamic Exp. 1 CE = -0.0015 ± 0.0004, t-Stat = -3.63, p < 0.001; Dynamic Exp. 2 CE = -0.0041 ± 0.0006, t-Stat = -6.46, p < 0.001; Dynamic Exp. 3 CE = -0.0059 ± 0.0011, t-Stat = -5.36, p < 0.001; Static Exp. CE = -0.0027 ± 0.0007, t-Stat = -4.02, p < 0.001).* *GLME model: ln RT ~ |TA-X_o_|<40° + (1|S); TA, target angle; S, subject. Exp.1 n = 12; Exp.2 n = 10; Exp.3 n = 10; Exp.4 n = 10. The error bars represent ±1 standard error of the mean.*
